# The rice leaf microbiome has a conserved community structure controlled by complex host-microbe interactions

**DOI:** 10.1101/615278

**Authors:** Veronica Roman-Reyna, Dale Pinili, Frances Nikki Borja, Ian Lorenzo Quibod, Simon C. Groen, Enung S Mulyaningsih, Agus Rachmat, Inez H. Slamet-Loedin, Nikolai Alexandrov, Ramil Mauleon, Ricardo Oliva

**Affiliations:** International Rice Research Institute, Manila, Philippines; Department of Biology, Center for Genomics and Systems Biology, New York University, New York, NY 10003, USA; Department of Plant Pathology, The Ohio State University, Columbus OH, USA. And Infectious Disease Institute, The Ohio State University, Columbus OH, USA; Research Center for Biotechnology, Indonesian Institute of Sciences. Cibinong, Bogor 16944, Indonesia

**Keywords:** *Oryza sativa*, Leaf microbiome, Abundance network, GWAS, functional profile

## Abstract

Understanding the factors that influence the outcome of crop interactions with microbes is key to managing crop diseases and improving yield. While the composition, structure and functional profile of crop microbial communities are shaped by complex interactions between the host, microbes and the environment, the relative contribution of each of these factors is mostly unknown. Here, we profiled the community composition of bacteria across leaves of 3,024 rice (*Oryza sativa*) accessions from field trials in China and the Philippines using metagenomics. Despite significant differences in diversity between environments, the structure and metabolic profiles of the microbiome appear to be conserved, suggesting that microbiomes converge onto core functions. Furthermore, co-occurrence analysis identified microbial hubs that regulate the network structure of the microbiome. We identified rice genomic regions controlling the abundance of these hubs, enriched for processes involved in stress responses and carbohydrate metabolism. We functionally validated the importance of these processes, finding that abundance of hub taxa was different in rice mutants with altered cellulose and salicylate accumulation, two major metabolites at the host-microbe interactions interface. By identifying key host genomic regions, host traits and hub microbes that govern microbiome composition, our study opens the door to designing future cropping systems.

## Introduction

Plant colonization of terrestrial habitats ignited the formation of biodiverse systems, termed phytobiomes, in which plants co-evolve with unicellular and multicellular organisms in fluctuating environmental conditions. In phytobiomes, plants are in constant interaction with microbial communities that adapted to colonize plant tissues, termed microbiomes (1). Microbes in these communities may have (nearly) neutral, harmful or beneficial effects on plant fitness. Benefits conferred by microbes to their host plants can be direct through protection from attacks and stressful environmental conditions, or indirect through the enhancement of plant resistance responses and/or plant growth (2–6).

Ecological theories suggest that microbiomes do not assemble randomly but that their formation is governed by complex interactions among microbes, host and environment (7,8). Understanding these complex interactions will help translational research to improve agronomic traits. The first step is to characterize and identify the mechanisms that drive the microbial community composition by quantifying the richness and diversity of taxa. The second step is to identify microbe-microbe metabolic interactions and host genetic factors, to define the ecological network structure (9–11). However, due to a lack of large-scale studies we still have only limited mechanistic insight into the identity and relative importance of factors that shape host-microbe interactions.

Asian rice (*Oryza sativa* L.) is grown globally and forms the staple food for over fifty percent of the world’s population. As part of the 3,000 Rice Genomes Project (3K-RGP) we recently completed the re-sequencing of a large genetic diversity panel comprised of 3,024 accessions from all major rice varietal groups(12,13). These accessions are adapted to a wide variety of agro-ecosystems and possess extensive heritable trait diversity, which in turn may influence microbiome assembly. Furthermore, the 3K-RGP panel has been used successfully to identify the genetic architecture underlying a number of complex morphological and phenological traits.

Here, we performed in-depth analyses of the meta-genomes of the 3,024 rice accessions to identify factors that drive microbiome assembly in the rice phyllosphere. We successfully captured the composition, structure, and functional profile of the leaf microbiomes of these accessions growing in two major areas of rice production, China and the Philippines. Our analyses showed that despite differences in the presence and abundance of individual microbial taxa the composition of the microbiome converges onto similar core metabolic functions in both environments. We discovered central taxa that as microbial “hubs” have an outsized influence on the network of host-microbe interactions and identified host genomic regions that control their abundance. These genomic regions were enriched for peroxisome-located processes involved in stress responses and carbohydrate metabolism. We functionally confirmed that host genetic variation in cellulose and salicylate accumulation can impact microbiome composition. The production of these compounds partially relies on peroxisome-located metabolism, and they play critical roles at the interface of host-microbe interactions. Our data provides insight into the mechanisms that drive microbiome assembly and opens the door for future initiatives to engineer consortia of beneficial microbes for crop performance improvement.

## Results

### Metagenome sequencing of the 3K-RGP accessions captures the rice leaf microbiome diversity

To characterize the rice leaf microbiome, we analyzed the metagenomic data from our 3K-RGP panel as explained in Fig. S1. Our panel contains sequencing data from 2,466 rice accessions grown in the Philippines (agPh) and 558 accessions grown in China (agCh). Microbial reads were identified after filtering against five reference rice genomes. Overall, 75% of the reads corresponded to Eubacteria and Archaea (Supplementary Table S1). We assessed species richness by measuring species accumulation (observed richness) and evenness. The accumulation curves across environments reached a plateau at 600 microbial genera present in a minimum of 100 rice accessions (Fig. S2A), and the average evenness values suggested a similar distribution of species abundance (Fig. S2B). Increasing the number of rice accessions in either agPh or agCh, did not result in the detection of more microbial genera. To account for differences in sample size, we calculated rarefied species richness and observed a higher number of genera in agCh (Fig. S2C). The total richness and alpha diversity (effective Shannon diversity) values were similar to other plant leaf microbiomes (11,14,15) and showed that agCh harbored 1.5 times more diversity than agPh (Fig. 1A-B; Supplementary Table S2). The higher values found in agCh might be associated with accessions being exposed to an array of microbial taxa missing or having lower abundance in the agPh environment (Fig 1C). Indeed, a major driver of microbiome diversity is the availability of microbes captured from the environment (9,10,16). Overall, our 3K-RGP meta-genomic sequencing effort successfully captured the leaf microbiome and identified the environment as a major factor that impacts microbial community diversity (9,17–20).

**Fig. 1.**
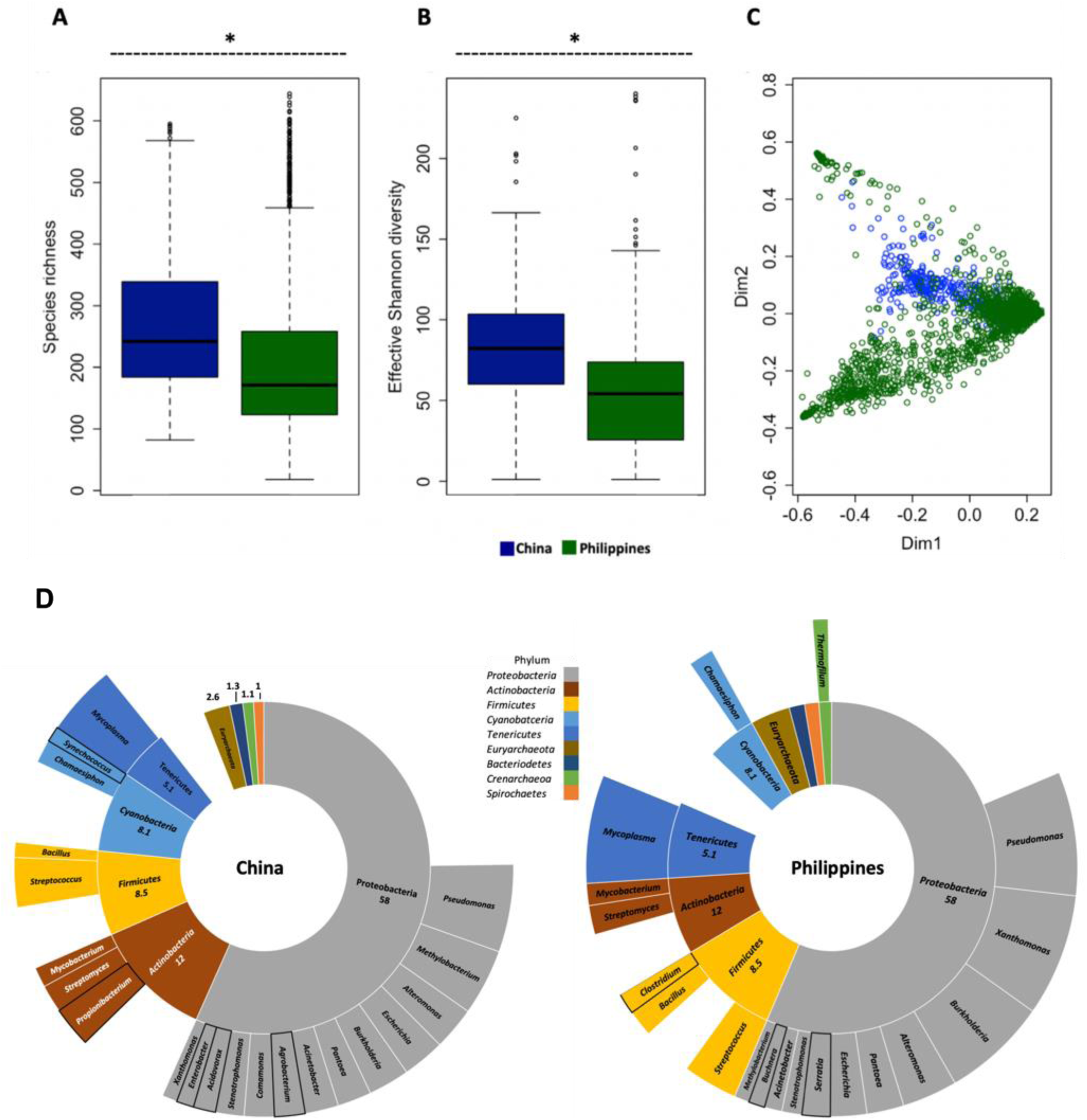
Host environment shapes the rice leaf microbiome diversity and composition. **A-B** The species richness and Shannon effective number of species comparisons between accessions grown in China and Philippines; *P-value < 0.001. Kruskal-Wallis test. **C** Weighted principal coordinates analysis based on the distances between environments microbial composition. The clustering is based on Bray–Curtis dissimilarity index. **D** Leaf microbiome composition of rice accessions grown in China and Philippines. The inner position of the sunburst chart represents taxonomic hierarchy Phylum and the outer position represents Genus. The chart shows abundance higher than 1%. The black line highlights the unique genera for each environment.

### Host environment and genotype shape the rice leaf microbiome

To further dissect differences in microbial community composition of agPh and agCh, we compared the relative abundance of Eubacteria and Archaea at different taxonomical levels. We found that 27% and 57% of the dissimilarity occurred at Phylum and genus level, respectively (Supplementary Table S3, S4). *Proteobacteria, Firmicutes, Actinobacteria, Cyanobacteria, Tenericutes*, and *Euryarchaeota* were the most abundant phyla (Fig 1D), resembling the abundances of these phyla in the leaf microbiomes of other crops (11,21–23). Interestingly, *Euryarchaeota*, which include methanogenic bacteria, and are frequently found under the anaerobic conditions in the rice paddies, were only marginally present in the aerobic phyllosphere (9,22). Presumably because of the aerobic conditions in the phyllosphere, we did not detect members of taxa commonly found in the soil or rhizosphere either (9,17–20).

While the microbiomes of agCh and agPh harbored 152 and 121 genera with relative abundance higher than 0.1%, respectively, only 25 genera contributed to the dissimilarities between them (Fig. 1D, Supplementary Table S3). These genera are common members of the leaf microbiomes of other crop plant species (9,24). For example, the genera *Propionibacterium, Agrobacterium, Acidovorax*, and *Enterobacter* were over four times more abundant in agCh than in agPh while *Xanthomonas* and *Serratia* showed the reverse pattern (Fig. 1D, Supplementary Table S4). Most of these genera include species that are oxygen-tolerant and capable of colonizing plant or animal hosts (Supplementary Table S4), contributing to a picture of the rice phyllosphere as a favorable environment for aerobic taxa. One factor that may account for the dissimilarities in the microbiome compositions of agPh and agCh, might be a difference in agricultural practices between the two environments. Different human interventions could lead to alternative routes in the horizontal acquisition of taxa in the microbiome (8,25,26). To confirm that agricultural practices could form a factor that shapes the leaf microbiome we needed to rule out that major genera were not artificially introduced during sample collection. To this end, we used qPCR to detect 11 highly abundant genera in 18 randomly selected accessions from our 3K-RGP panel (Supplementary Table S5). We were able to quantify the presence of all taxa and observed a similar distribution across accessions (Fig. S3). Similar to our previous findings, the genera *Pseudomonas*, *Xanthomonas*, *Mycoplasma*, and *Mycobacterium* were the most abundant genera, ruling out that highly abundant genera were introduced artificially.

Another factor that shapes the microbiome is the host genotype (10,16,19,27). To evaluate the extent to which host genotype impacts the assembly of microbial communities, we compared a number of diversity indices among 12 rice varietal groups and proximal clusters that act as a proxy for host genotype (13). We found that the richness and evenness values were strongly influenced by host genotype, whereas the diversity value was affected by the environment (Fig. 2, Supplementary Table S6). Eight out of 12 rice varietal groups and proximal clusters harbored a similar number and distribution of microbial genera, independent of the fact that plants in these groups and clusters were grown in different environments. Despite this significant effect of host genotype, heritability values for the most abundant genera in agPh and agCh were relatively low (Fig. S4) (16,23). This pattern would be expected if accessions in the groups and clusters each carry different combinations of alleles underlying trait differences that influence microbiome composition, i.e. if the genetic basis for such traits is diffuse. If this is the case, then certain host traits should explain more of the variation in the composition of the rice leaf microbiome.

**Fig.2.**
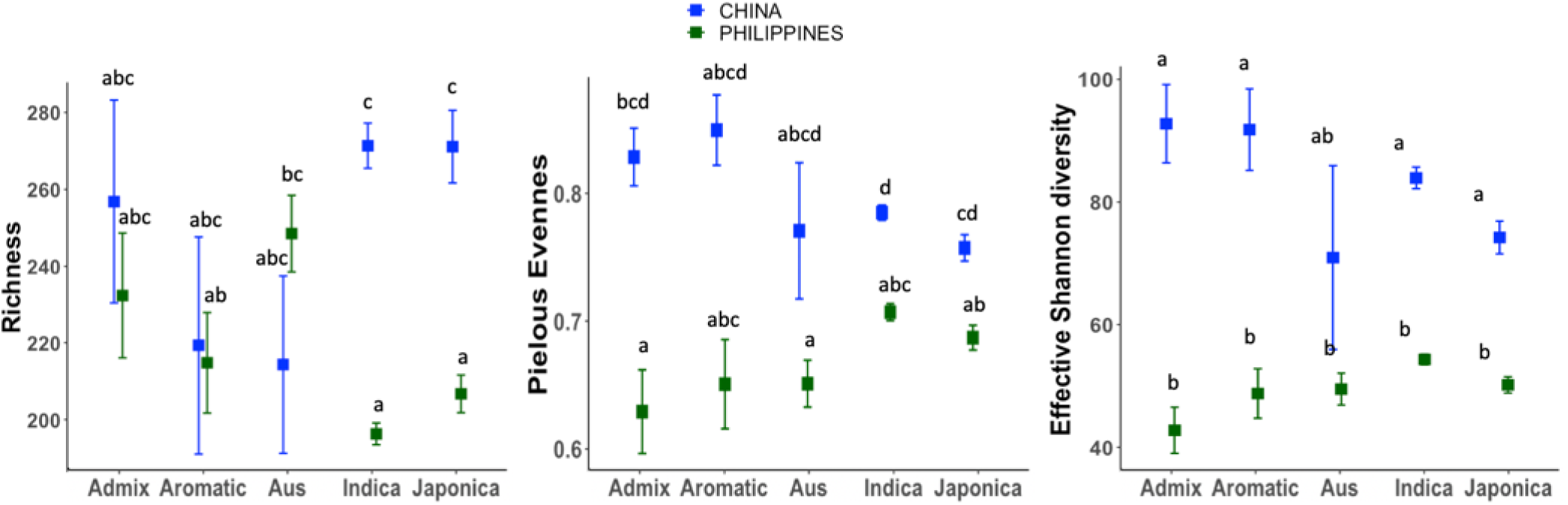
Rice varietal groups and the environment shapes the leaf microbiome. Least squares mean estimates of leaf microbiome richness (left panel), evenness (middle panel) and effective Shannon diversity (right panel) in the rice varietal groups (Admix, Aromatic, Aus, Indica and Japonica) after environment adjustment. Means sharing the same letter are not significant different based on Tukey method (alpha = 0.05). The analysis for all varietal groups is in the Supplementary Table S6.

Accessions in different rice varietal groups and proximal clusters are known to have adapted independently to the same agro-ecosystems, often converging on the same traits. Indeed, we find that the agro-ecosystem in which accessions were originally collected explained a significant amount of variation in microbiome composition independent of the experimental conditions (Fig. S5, Supplementary Table S7). Overall, our data suggest that environment plays a key role in determining variation in microbial community composition. However, other factors such as host genetic background and traits associated with adaptation to particular ecologies further condition the assembly of the leaf microbiome (28–30).

### The rice leaf microbiome structure is conserved despite differences in community composition

The establishment and maintenance of the microbial community is further shaped by networks of interactions among microbes (1,31,32). To identify essential microbial relationships, we inferred co-occurrence ecological networks from the agCh and agPh datasets. Interestingly and despite significant differences in composition, agCh and agPh assembled communities with similar structures (Fig. 3A), suggesting similar network properties. Although the mechanism that favors co-existence of highly diverse microbial community remains uncharacterized, this shows that community assembly might follow specific rules independent of the availability of taxa for recruitment. We identified seven highly connected genera or “hubs” in networks of microbes colonizing agCh and agPh (Fig. 3A, Supplementary Table S8-S9). Moreover, the networks for plants in both environments shared *Clostridium*, *Mycoplasma*, and *Helicobacter* as three hubs with the highest number of connections and positive associations (r_Pearson_ > 0.7, P-value < 0.001) (Supplementary Table S7, Supplementary Table S8). The hubs genera appear to stabilize the network of interactions because when we artificially removed these genera from the analysis the interactions were lost (Fig. S6). Similar to other studies (31,33), our results suggest that the hub genera have a regulatory effect on the network of microbial interactions and or may play an important ecological role in the microbial community. The connectivity of a genus within the network did not correlate with the abundance of that genus. For example, the highly-abundant genera *Xanthomonas* and *Streptococcus* were not identified as hubs, while *Helicobacter*, with less than 1% abundance, still plays a role in shaping the network of interactions (33–35). We are aware that other inter-kingdom interactions might be driving the differences in microbial community composition (33,36), but their influence appears to be limited in the case of the rice leaf microbiome.

**Fig.3.**
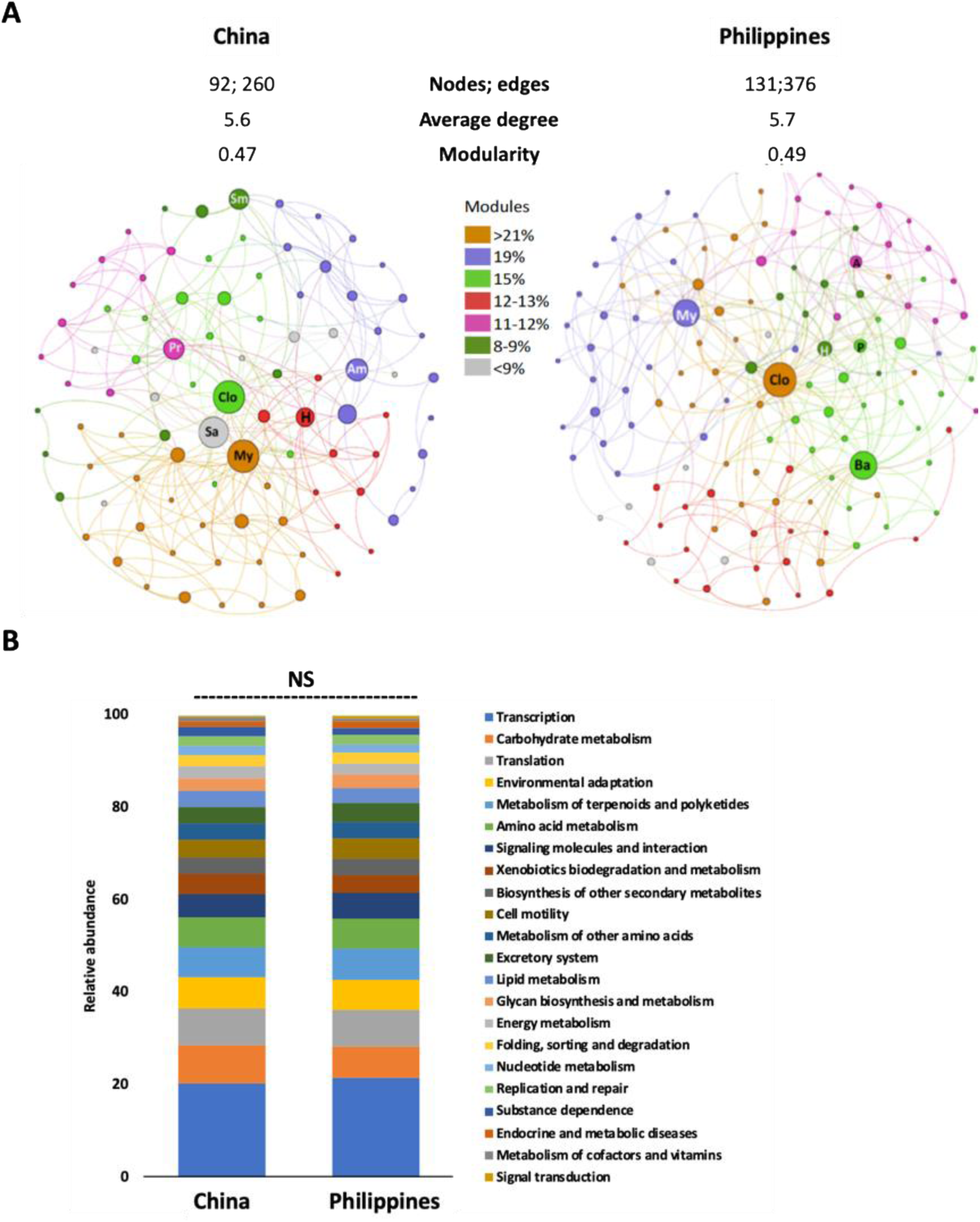
Microbial ecological network of the rice leaf microbiome identified common hubs that could explain the conserved functional profile among environments. **A** Microbial ecological network from China and Philippines with abundant genera present in at least 50% of all samples. The colors represent the seven modules of each network. Each node represents a genus and the circle size indicates betweenness centrality increment. The key microbial hubs are *Clostridium (Clo), Mycoplasma (My) and Helicobacter (H)*. Other hubs in China are *Spiroplasma (Sa), Azospirillum (Am), Prochlorococcus (Pr), Sphingobium (Sm)*. For Philippines, important hubs are *Bacillus (Ba), Pseudomonas (P)*, and *Azotobacter* (A). The properties of the network are number of edges, number of nodes or genera, average degree and modularity. Only for the network analysis the genus counts were center-log-transformed. **B** KEGG level 2 pathways with more than 1% relative abundance in accessions grown in China and Philippines. NS no significant, Wilcoxon rank-sum test = 6869, P-value = 0.421.

We next analyzed the network topology to reveal modular interaction patterns. We found that the networks for both agCh and agPh had seven modules (Fig. 3A, Supplementary Table S8-S9). This structure suggests a highly stable network since a microbial community appears to reach an equilibrium when its network of interactions has a small number of modules (31). Compared to other studies (9,20), we found that most of the modules were not organized randomly but rather shaped by microbial genetic ancestry, biological function, or ecological niche. For instance, we found that one module was enriched with Cyanobacteria, while another module showed enrichment for plant growth-promoting bacteria (Supplementary Table S7 and S8). This aligns with the idea that some microbes in the phyllosphere adapted to the leaf surface conditions (31). While the data suggest that members of the same module might have strong biochemical associations, it is not clear if modules overlap in the roles they play, or if members co-localize to the same leaf sub-compartments. Overall, the leaf microbiome structure is likely defined more strongly by the biological functions of modules than by the diversity or abundance of individual taxa (23,37–39). The fact that we identified the same hubs genera in two independent datasets, from two different rice growing environments, strongly aligns with the ideas that hubs have an outsized role in shaping the microbial community and that community assembly follows certain rules (1).

### The functional profile of the leaf microbiome is conserved despite differences in microbial composition

If the structure of the microbial community is defined by rules that depend on the overall function of each module of microbial taxa (1,31,34), then the microbes of agCh and agPh should share metabolic profiles. We predicted functional categories for the microbial taxa and found that the communities on agCh and agPh exhibited similar profiles (Fig 3B, Supplementary Table S10). Both datasets shared 22 of 24 KEGG level 2 pathways (Fig 3B, Supplementary Table S10). The most abundant pathways, also common to other leaf microbiomes, were associated with transcription, carbohydrate metabolism, translation, environmental adaptation, metabolism of terpenoids, and amino acid metabolism (20,37,40,41). The presence of common categories such as xenobiotic biodegradation and biosynthesis of secondary metabolites might indicate microbial adaptation to agricultural practices in each rice-growing environment (1,22,42). Interestingly, the microbes of agCh contained five more pathways for carbon fixation, than those of agPh, which might be caused by adaptation to differences in climatic conditions or light exposure on the leaf surface (43,44). The redundancy of functions and hub genera in the leaf microbiomes of both environments supports the idea of a core set of microbes in each community that is under selective pressure from complex microbiome-host interactions to provide essential functions for the community, which aligns with the concept of a functional entity or holobiont (1,43).

The nature of our experiment prevented us to test if the leaf microbiome had different metabolic profiles compared to microbial communities in the soils or roots in the same experimental conditions in the two environments. Due to these limitations, we used other available plant microbiome datasets to estimate functional categories. We evaluated three shotgun metagenome sequencing datasets (rice soil, wheat soil, and rice leaf) and three 16S rDNA databases (rice root endosphere, rice rhizosphere, and maize leaf) (Supplementary Table S11). As expected, samples profile with different sequencing technologies formed separate clusters (Fig. S7). Despite the fact that 16S rDNA and shotgun meta-genome sequencing captured different types of information, samples profiled with either technique showed that microbiomes of leaves, roots, and soils have consistently different functional profiles (Fig. S7). This result aligned with other studies finding that biological functions of microbial community are linked to the plant tissues and soil compartments of origin (33,38,41,45). In summary, it is likely that key microbial genera (“hubs”) make an important contribution to organizing the leaf microbiome as a network of microbial modules that perform a complex of functional roles tailored to the phyllosphere.

### Rice metabolic pathways modulate the leaf microbial community

To identify rice genetic factors that may control the recruitment and establishment of the microbial hub genera, we conducted a genome-wide association study (GWAS) on 3,024 rice accessions, using the genomic information from 6.5 million SNPs and the relative abundance of the three key hubs that were present in both environments: *Helicobacter*, *Mycoplasma*, and *Clostridium*. Overall, we found 32 significant SNPs associated with hub abundance (P-value < 1E-15, ci = 0.95), distributed across nine chromosomes (Fig. 4A, Supplementary Table S12). Thirty out of 32 SNPs were located within 16 annotated rice genes (Supplementary Table S13). Eleven SNPs had a missense effect on nine of the 16 genes. Linkage disequilibrium analysis identified 19 haplotype blocks ranging from 30 to 130 kb and spanning 180 candidate genes (Supplementary Table S14).

**Fig.4.**
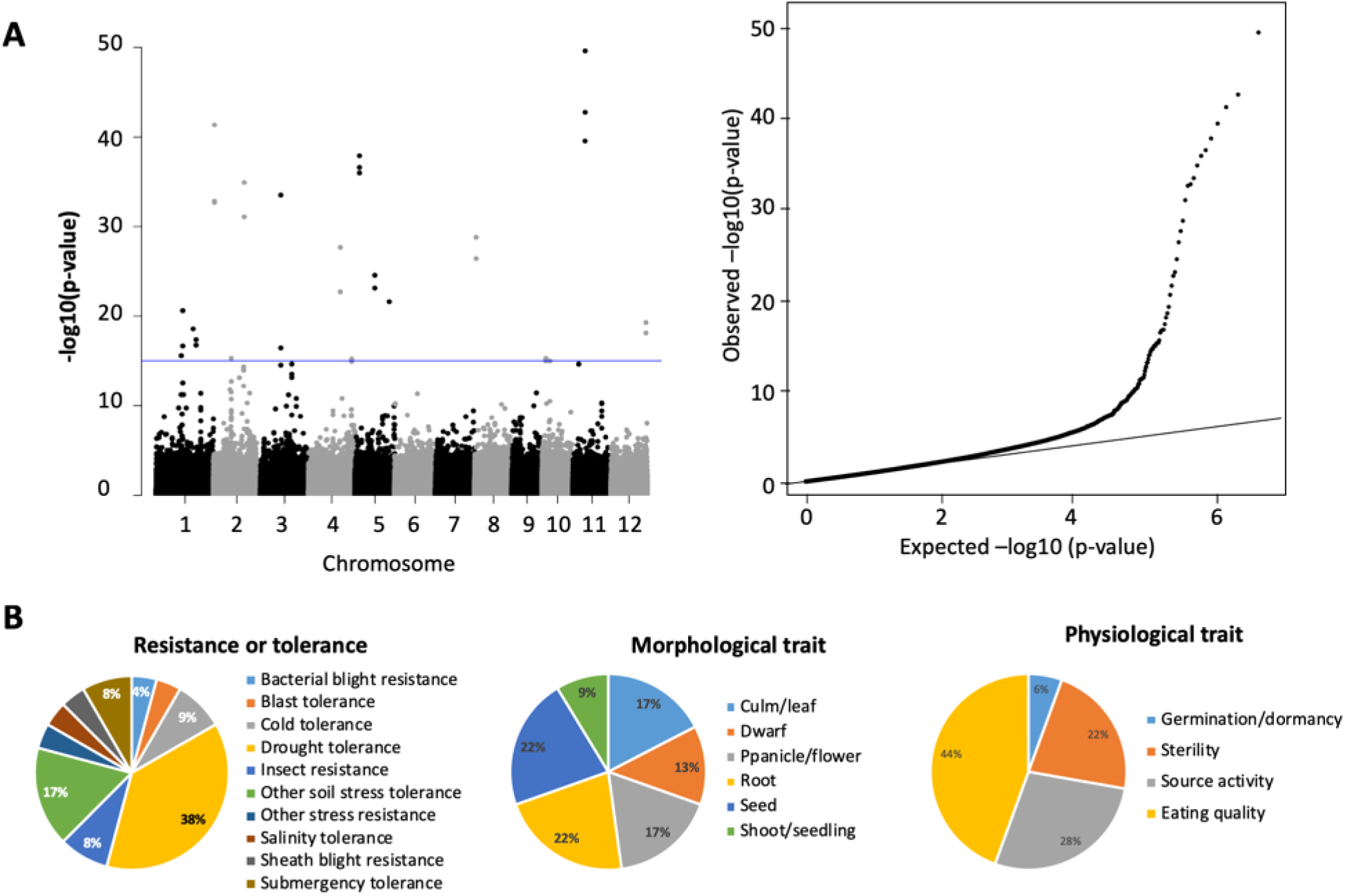
Rice metabolic pathways are associated with the microbiome structure. **A** Genome wide association study for the three microbial hubs in combine environments. Manhattan plot (left) and quartile–quartile plot (right) indicate major peaks (significant SNPs) associated with microbial abundance. P-values were adjusted with FDR and values lower than 1E-15 were consider significant (blue line). The significant hits are distributed across nine chromosomes. **B** The significant SNPs found in this study co-localize with a number of agronomic QTLs categorized as: resistance or tolerance, morphological trait and physiological trait. Categories were retrieved from Q-Taro database.

To assess if any of these genes have been previously associated with rice agronomic traits, we used the rice quantitative trait locus (QTL) database, Q-TARO. Overall, the 180 candidate genes mapped to 65 QTLs distributed in different categories: biotic or abiotic stresses (24 QTLs), morphological traits (23 QTLs), and physiological traits (18 QTLs) (Fig. 4B, Supplementary Table S14). In addition, 36 candidate genes were connected to each other in pathways related to stress responses, carbohydrates metabolism, and amino acid metabolism whose enzymatic reactions take place for an important part in the peroxisome (Supplementary Table S15). Host genetic factors involved in the same processes were linked to leaf microbiome assembly in similar studies in Arabidopsis, Nicotiana, and maize (23,32,46,47). This makes it likely that allelic variation in certain host genes influences he abundance of hub genera and this the composition of the rice leaf microbiomes.

To test the hypothesis that single host genes involved in stress responses and carbohydrate metabolism could modulate the leaf microbiome, we performed 16S rDNA sequencing on the apoplastic fluid of rice lines with different accumulation of salicylate and cellulose – two compounds whose production takes place partially in the peroxisome. For cellulose accumulation, we compared microbial community profiles on the Indica rice cultivar IR24 and its Xa4-containing near-isogenic line (IR24+Xa4). The protein XA4 is a cell-wall associated kinase involved in cellulose accumulation, which influences leaf mechanical strength and defense responses to bacterial infection (48). For salicylate production, we compared the microbiomes of the Japonica rice line Rojolele (accession R711) to its overexpressing line (R711+SAox). The latter line constitutively expresses the bacterial genes entC and pmsB, encoding for isochorismate synthase and isochorismate-pyruvate lyase, respectively (49). Both genes are involved in the salicylic acid biosynthetic pathway, a phytohormone with a key role in stress responses. In both scenarios, the bacterial richness decreased, while the abundance of *Proteobacteria, Firmicutes, Actinobacteria*, and *Bacteriodetes* fluctuated when the alleles of interest were present (Fig. 5 A-D, Supplementary Table S16). The line IR24+Xa4 showed a reduction in the abundance of *Actinobacteria*, but an increase in *Proteobacteria* and *Firmicutes* (Fig 5A). At the same time, all phyla showed a substantial decrease in the number of genera by which they are represented in the microbiome (Fig. 5C). The line R711+SAox had a decrease in the abundance of *Firmicutes* and increase in *Proteobacteria* (Fig 5B). Nevertheless, the reduction in the number of genera present was less dramatic than in the IR24 – IR24+Xa4 comparison (Fig. 5C, D). The small difference detected in abundance, despite the decrease in the number of genera present, indicated that the remaining genera acclimated or adapted to the stressful environmental presence of increase salicylate accumulation by occupying the spaces left by the decline of other genera. A closer look at the hub genera showed that the majority of hubs remained unaltered, and that only two hub genera experience significant changes in abundance between the lines (Fig. 5E, F). For instance, *Bacillus*, *Pseudomonas*, and *Helicobacter* increased in abundance in IR24+Xa4 and R711+SAox lines compared to IR24 and R711, respectively. In the case of *Clostridium*, the salicylate and cellulose accumulation appears to correlate with a reduction in the abundance of this hub. Finally, *Sphingobium* abundance change was specific to the stress signal. In IR24+Xa4 the abundance of this genera increased while in R711+SAox decrease, compared to their respective controls. The exacerbation of cellulose and salicylate accumulation in these lines appears to modulate the presence of specific microbial groups in the apoplast, suggesting that the host might reshape the composition of the microbial community in a controlled fashion. Additional evidence is needed to understand the driving forces behind the modulation of the abundance of hub genera and any independent or knock-on effects on the abundance of other microbial taxa.

**Fig. 5.**
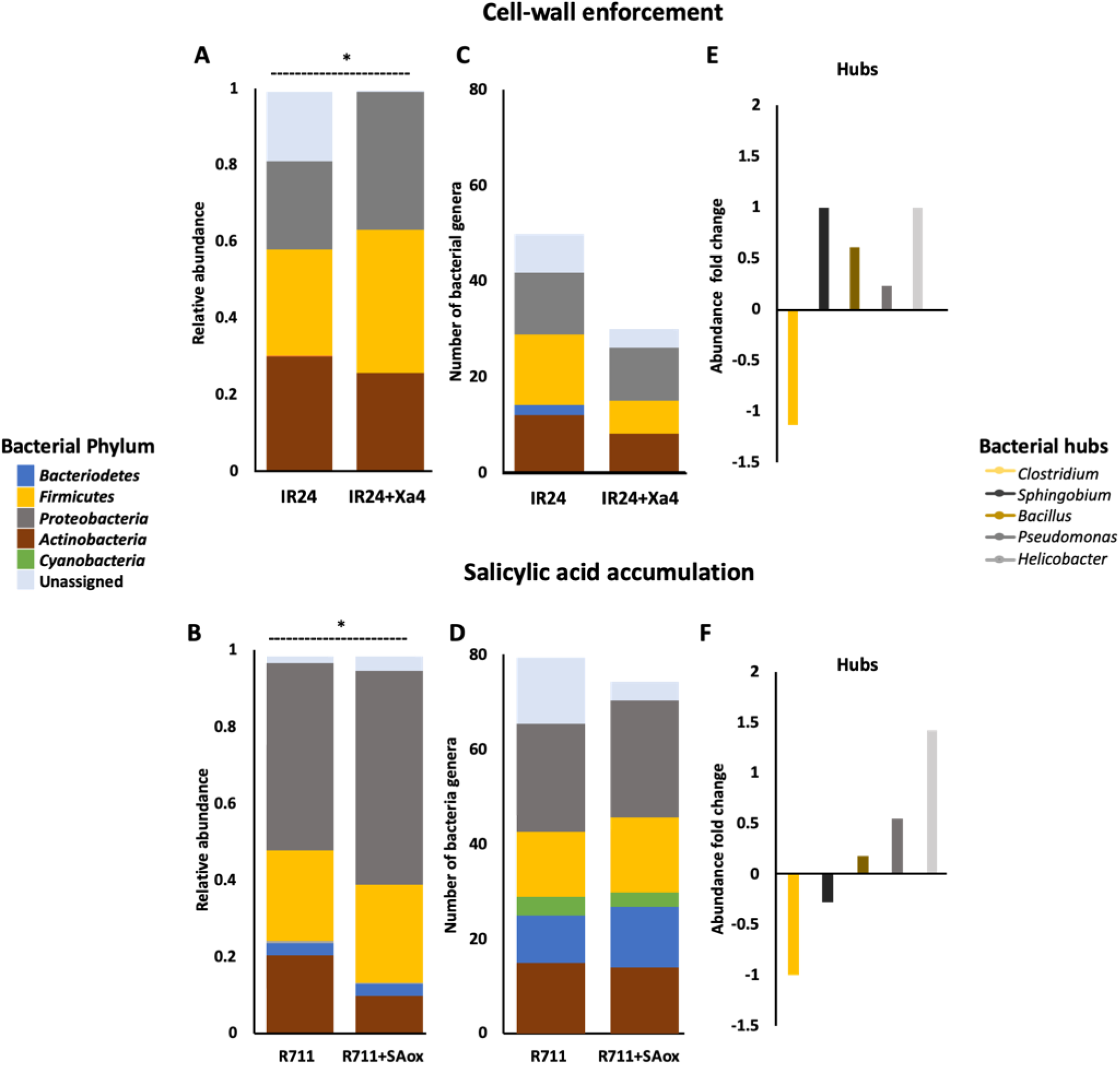
Host genes related to stress responses modify microbiome composition. **A-B** Phyla-level distribution in rice lines with different accumulation of cellulose (IR24+Xa4) and salicylate (R711+SAox) based on 16S rDNA amplicon. *P-value <0.05, Wilcoxon rank-sum test. **C-D** Genera-level numbers for each Phylum in each rice line. **E, F** Microbial hubs abundance fold change between control and rice line with modified accumulation.

### Conclusions

Microbial communities that live in association with plants carry a great diversity of metabolic capabilities and often influence broad aspects of plant biology. In agricultural environments, the composition of these communities affects overall crop performance by contributing to important plant functions such as vegetative growth, nutrient uptake, and immune responses, among others (2,6,41). Efforts to understand and exploit such capabilities may bring exciting opportunities to design future cropping systems. Using meta-genomic profiling of the 3K-RGP panel, we described the regulatory factors that shape the rice leaf microbiome. Our results indicated that the environment is the main reservoir of microbial diversity. Common agricultural practices, such as crop irrigation or the use of animal labor, might also explain how microbes from other niches are usually part of the phyllosphere. The structure of the leaf microbiome is most likely determined by ecological networks that perform core functions. Some of these functions, such as carbon fixation or xenobiotics degradation, suggest adaption to the leaf environment in the context of modern agriculture. Moreover, the networks revealed key microbial groups that regulate the establishment of the community but also appear to be controlled genetically by the host. It is not surprising that some of the identified regions are enriched in genes related to stress response since the microbiome evolved to interface and react to environmental variation. Our results validate the idea that both, the plant and the microbiome, shape the network of interactions and therefore co-evolutionary tracks are inevitable. Give the scale of the dataset, we have taken the first steps in unearthing the factors behind microbiome assembly in rice, which can be harnessed for engineering future crop improvements.

## Methods

### Genomic source

To describe the rice leaf microbiome, we used the 3,000 Rice Genomes Project database (12). This database was originally created to gather information about the rice genetic variation. The lines were planted in two different environments. Around 2,466 accessions at the International Rice Research Institute, in the Philippines, and 558 accessions at the Chinese academy of Agricultural Science in China. The lines include five varietal groups: Indica, Japonica, Aus, Aromatic and admixed (13). Indica and Japonica can be further subdivided into genetically proximal clusters (13). Japonica has four clusters (Japx, Tropical Japonica (named trop), Subtropical Japonica (named subtrop), and Temperate Japonica (named temp)). Indica has five clusters (Indx, Ind1A, Ind1B, Ind2, and Ind3). The database also includes information on country of origin, breeding classification, and ecosystem. Here we repurposed the database to gather information about the rice leaf microbiome (see Figure S1 for details about the project). We mapped each rice accession genome to the five reference rice genomes (Nipponbare, 93-11, IR64, Kasalath, and DJ123) with the software BWA v0.7.10 (50). We extracted the reads that did not map to the five rice genomes with samtools v1.0 (51). The reads were converted to Fasta files with BEDtools v2.17.0 (52) and used as entries for the software Kraken v1.0 (53). This software classified the reads from Phylum to Genus-levels based on the bacteria and archaea database from RefSeq NCBI database (release 69). To estimate taxa abundance we used the Bayesian-based tool Bracken v1.0 (54). We kept the genera that were present in at least 10% of samples for further analysis.

### Diversity estimation

For composition analysis we used the relative abundance normalization on the count matrix, where the read counts for a taxa-level in a given sample were divided by the sum of all counts in that sample. To calculate the richness and diversity indexes, we use the R package Vegan v2.5-3. To check homogeneity of variance across samples we used the classical Levene’s test with mean. Comparison of alpha diversity values were performed with ANOVA and the linear model y ~ environment, where y is the richness, evenness or effective Shannon diversity. For ad-hoc analysis we used Wilcoxon and Kruskal-Wallis tests. To calculate dissimilarity indices in the microbial community, we run the Vegan function vedgist with the Bray-Curtis method and the function wcmdscale to plot a weighted principal coordinates analysis. To identify which taxa contributes to dissimilarities between environments, we used the function Simper from the R package Vegan. We used the relative abundance of Phyla or genus as microbial community matrix, environment as grouping factor and 100 permutations. For comparison among the rice varietal groups and among clusters, we set the linear model y ~ rice varietal group*environment, where y was richness, evenness or effective Shannon diversity. Then we adjusted the linear model to the means of the factor environment with the least-squares means function in R package Emmeans v1.3.3. We used 95% confidence interval and Tukey-adjusted comparisons. To estimate broad heritability of the most abundant genera within environment, we used the R package lme4 v1.1-19 to fit a random effect linear model. The abundant genera were normalized to relative abundance and the fixed variable was rice varietal group. To estimated heritability, we divided the variance of the model to the sum of all variances and residuals. We plotted the values for each environment and genera. To evaluate if other factors shape the microbial community, we used 467 accession grown in Philippines that have full information about country of origin, breeding classification, and ecosystem. We calculated a distance matrix with the R package Vegan and visualized the distribution of microbial abundance taxa with a canonical correspondence analysis. For correlations we used the chi-square values.

### Quantification of 16S from abundant genera

To validate the results from 3K-RGP metagenome analysis, we amplified and quantified eleven of the most abundant genera in 18 randomly selected rice accessions from the 3K-RGP. The 18 accessions were five Indica, five Japonica, two Aus, four Admix, and two Aromatic. We grew the plants in glasshouse conditions at The International Rice Research Institute and harvested the leaves at 21 days old. We cleaned the leaves with ethanol, bleach and water before DNA extraction. DNA was extracted with CTAB method (12). The DNA was aliquoted in similar concentrations for the qPCR. For amplification and quantification, we used the StepOnePlus™ Real-Time PCR System and SYBR Green following manufacturer protocol (Applied Biosystems). We selected published primers for *Pseudomonas* sp., *Burkholderia* sp., *Mycoplasma* sp*., Streptomyces* sp., *Methylobacterium* sp and 16s rDNA region V34 (Supplementary Table S6). We designed primers for *Mycobacterium* sp., *Xanthomonas sp., Alteromonas sp., Pantoea sp., Spiroplasma* sp., *Bacillus sp. and Clostridium sp*. For comparisons, all samples were normalized to the 16S rDNA region V34 and plotted in logarithmic scale. We included primers for *Spiroplasma* to validate that the reads assigned to *Mycoplasma* were not a wrong annotation.

### Microbial ecological network and functional analysis

For the microbiome microbial ecological network we used the program SpiecEasi v0.1.4 (55). As the program is sensitive to rare species, we removed all the genera that were not present in at least 50% of all the samples from the count matrix. We used the absolute counts as the program does a center-log-transformation. We did the analysis with the Meinshausen-Buhlmann’s neighborhood method and the following parameters, lambda.min.ratio=1e-2, nlambda=20, pulsar.params=list (rep.num=100, ncores=7) (55). The program was run in R and the network was plotted with Gephi v0.9.2. The functional profile for agCh and agPh was predicted with the web-based tool Vikodak v1.0 under the co-metabolism algorithm workflow (56). Briefly, this algorithm is based on the assumption that genes present by various microbes in the microbial community contribute to specific metabolic pathway(s). The functions were classified with KEGG hierarchy levels. For further comparisons, we kept pathways with more than 1% abundance. We run a Wilcoxon rank sum test to compare the agCh and agPh microbial profiles.

### Genome Wide Association Study

We implemented PLINK 1.9 (61) and GEMMA 0.97 (58) for the population stratification and SNP-based association test. We kept Chinese (agCh) and Philippines (agPh) rice accessions together because some agCh lack SNP information, which will bias the association analysis. For the analysis we used 6.5 million filtered rice SNPs from the 29 million bi-allelic SNPs retrieved from the Rice SNP-Seek Database v0.4 (snp-seek.irri.org). We excluded SNPs with lower genotypic rate (>95%) and minor allele frequency (MAF < 0.01). We removed the SNPs that fail the Hardy-Weinberg equilibrium test (P < 0.0001). We performed GWAS with the centered log ratio-transformed abundance of the three hubs *Clostridium, Helicobacter* and *Mycoplasma* as phenotypic trait. We used abundance because our hypothesis is that hubs have a strong effect on the microbiome interactions, and the ecological network was build based on co-abundance. We also run the analysis using other genus from the network and we found overlapping in some SNPs. We run GWAS with the GEMMA multivariable linear model and identified significant SNPs by filtering with False Discovery Rate (FDR <0.01) and P-value (P-value <1E-15). The Manhattan plots and quantile-quantile (Q-Q) plot were created with the R package qqman v.0.1.3. We determine the expected and observed probabilities of SNPs association with Q-Qplot. We grouped the significant markers by haploblocks based on the linkage disequilibrium decay (LD<0.3) and correlation coefficients (*r^2^* > 0.6) in each chromosome using Haploview v4.2 (59,60). We identify and described the genes by gene ontology annotation, QTL overlapping, RiceNet v2 interactions and SNP effect based on the information from SNP-Seek Database (snp-seek.irri.org) (Dataset S13).

To validate the GWAS results, we evaluated the effect of stress response pathway on the microbiome composition. We selected rice lines available at the International Rice Research Institute. The rice lines IR24 and R711 were used as controls and compared to the lines IR24+Xa4 and R711+SAox which had altered cellulose and salicylate accumulation levels, respectively. The rice line IR24+Xa4, has the gene Xa4 in the IR24 background. The gene Xa4, is associated with cell-wall reinforcement and was introduced as part of breeding programs. The line R711+SAox has the genes entC and pmsB, related to the salicylic acid biosynthetic pathway. The gene construct containing both EntC and pmsB genes under CaMV 35S promoter fused with plastid targeting sequence (49) was inserted to a modified pCAMBIA 1300 and transformed in the rice cultivar Rojolele accession number R711 following the modified method of Toki et al (61). The presence of transgenes in the progenies were detected by PCR amplification. We extracted the leaf apoplastic fluids from IR24, IR24+Xa4, R711 and R711+SAox and recover the 16S rDNA by PCR amplification. We used apoplastic fluids instead of whole tissue to avoid overrepresentation of plastid DNA and to reduce noise by using only endophytes. Using apoplast, instead of whole tissue, we reduced 80% the chloroplast contamination. For the apoplastic fluids extraction, we used negative pressure with a syringe to force water into the apoplast and then by centrifugation (1000 rcf, 10 min, 4 C) wash out the apoplastic fluids. For the 16S enrichment we performed a PCR with Q5® High-Fidelity DNA Polymerase (New England Biolabs), the forward primer 341F and the reverse primer 806R to cover the V3/V4 region. To test for bacteria contamination, we did a PCR with the water used for apoplast extraction. If we did not observe bands with water as template, we pooled PCR products from six samples of the same rice line and send the pool for sequencing. We sent 5 ug (total mass) of pooled PCR products to BGI group (https://www.bgi.com) for 16S Amplicon Sequencing with Illumina MiSeq PE300 using the 16S V3-V4 region. BGI gave us, on average, 125,000 cleaned paired end reads of 300 base pairs. We confirm the reads were clean using the programs Trimmomatic v0.38 (SLIDINGWINDOW:5:15 MINLEN:200 AVGQUAL:20) and Flash2 v2.2.00 (-m 10 -x 0.1 -M 200) (62,63). The downstream analysis were done with the Qiime2 v 2018.11 and the “moving pictures” tutorial (64). Briefly we used dada2 to detect and correct Illumina amplicon sequence data. We assigned taxonomy to the sequences using the Small Subunit (SSU) rRNA Database from Silva release 132 (https://www.arb-silva.de/).

## Author contributions

NA, RO, designed research; VR-R, DL, FNB, ILQ, RM, IS-L, ESM, AR performed research; VR-R, DL, RM, RO analyzed data; VR-R, SCG, RO wrote the paper.

## Acknowledgements

Authors would like to thank Oscar Lujan (Northern Arizona University) for process the samples, the Rice CRP and Department of Science and Technology – Advanced Science and Technology Institute (DOST-ASTI) for the free access of high-performance computing services. DP was funded by the DOST-ASTHRDP program. VRR was funded by the Newton Fund. S.C.G. was supported by a grant from the Gordon and Betty Moore Foundation (Life Sciences Research Foundation Postdoctoral Fellowship Grant GBMF2550.06 to S.C.G.)

## Captions for supplementary tables

**Supplementary table S1.** List of 3K-RGP rice accessions with number of reads that did not map to the rice genomes (unmapped reads). The reads were subjected to taxonomic classification with the software Kraken (Kraken output) and were quantified with the software Bracken to Phylum and Genera (Bracken output). The growing location for each accession is also listed.

**Supplementary table S2.** List of 3K-RGP rice accessions with the richness and diversity indexes. The indexes were total read counts (same bracken output), the logarithm of total counts (Log10_TRC), the number of genera (Genus_Counts), Effective Shannon diveristy, Pielou’s evenness and Simpson 1/D index.

**Supplementary table S3.** Significant Phylum and Genera that contribute to the differences between accessions grown in China and accessions grown in Philippines. The biology of the 25 genera is indicated as tolerance to oxygen and niche.

**Supplementary table S4**. Relative abundance (average and standard deviation) of the 533 genera in accessions grown in China and accessions grown in Philippines.

**Supplementary table S5.** Sequences of 16S primers used for validation of metagenomic analysis and list of the 18 rice accessions from the 3K-RGP project used for validation. The primers were based on other publications or design for this study. The primers that amplify the rice actin gene were used as control.

**Supplementary table S6.** Least squares mean estimates of leaf microbiome richness, evenness and effective Shannon diversity in the rice varietal groups (Admix, Aromatic, Aus, Indica and Japonica) and clusters.

**Supplementary table S7.** Description for 2,234 lines with reliable passport data from the IRRI database or accession grown in Philippines.

**Supplementary table S8.** Co-abundance network values for the most abundant genera in accessions grown in Philippines.

**Supplementary table S9.** Co-abundance network values for the most abundant genera in accessions grown in Philippines China.

**Supplementary table S10.** Metabolic pathways predicted by Vikodak for each environment, based on KEGG levels 1,2 and 3. Average and

**Supplementary table S11.** List of NCBI microbiome accessions used for the correspondence analyses on the functional profiles of 16S and shotgun sequencing technologies.

**Supplementary table S12:** Significant signals from the genomic wide association analysis (GWAS) with a multivariable linear model using 6.5 million SNPs and the three hubs abundance. We kept SNPs with a P-wald value lower than 1E-15.

**Supplementary table S13:** Description of the significant SNPs, from Supplementary table S11. Chromosome, genomic position in Nipponbare genome, annotation, gene ontology and SNP effect were retrieved from the webpage snpseek.org.

**Supplementary table S14:** Description of haplotype blocks for each significant SNP, number of associated candidate genes and the QTLs that match to the same region.

**Supplementary table S15:** Interactions between all the candidate genes associated with the haplotype blocks. The analysis was retrieved from RiceNet webpage. The probabilistic functional network database for interactions was AUC= 0.92; P-value < 0.0001.

**Supplementary table S16:** Relative abundance of the apoplastic microbiome from IR24, IR24+Xa4, R711 and R711+SAox. The confidence value indicates the average classification of reads to that group.

## Supplementary figures

**Figure S1.**
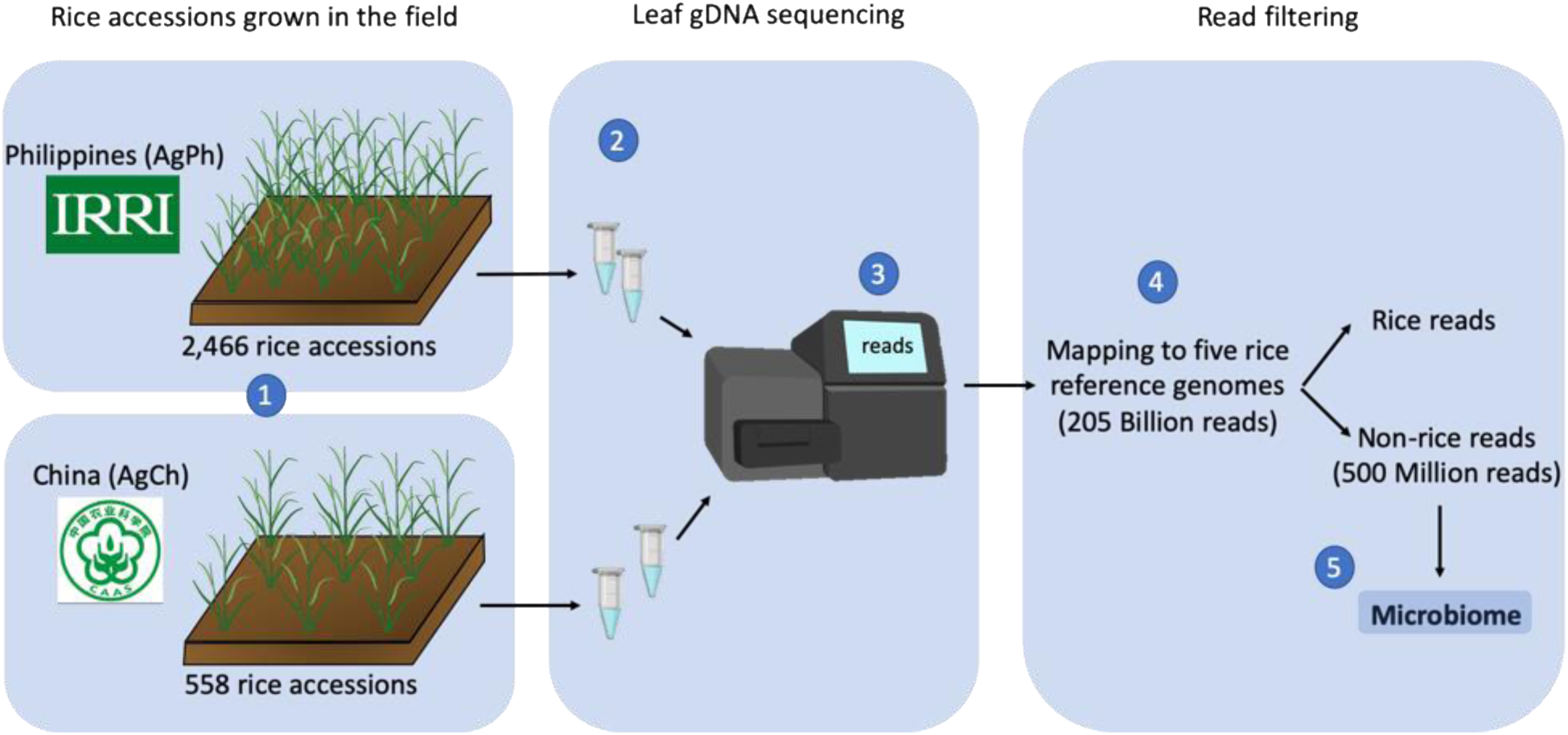
Generation of 3000 rice genomes dataset and pipeline for collecting the leaf microbiome. **1.** Selected gene bank accessions where grown at Philippines (agPh) or China (agCh). The Philippines accessions belong to the International Rice Gene bank Collection (IRGC) at the International Rice Research Institute (IRRI). The accessions grown in China are part of a bigger collection from the China National Crop Gene Bank (CNCGB) in the Institute of Crop Sciences, Chinese Academy of Agricultural Sciences (CAAS). The rice accessions were grown in the field and the environmental conditions between China and Philippines were more likely different (12). **2.** Genomic DNA (gDNA) was extracted from young leaves of each accession by modified CTAB method. **3.** All genomes were sent to BGI group (https://www.bgi.com) to construct the libraries and do the sequencing with the HiSeq2000 platform. **4.** Clean reads, that correspond to 205,084,357,762 paired-end reads for all 3,024 genomes, were then map to five reference genomes using the BWA software. The reference genomes are Nipponbare, 93-11, IR64, Kasalath, and DJ123 (13). We separate the reads that map to all rice genomes from the reads that did not map to any rice genome. **5.** We suggest the reads that did not map to any of the rice genomes (non-rice reads) came from microbial DNA that cohabit with rice.

**Fig S2.**
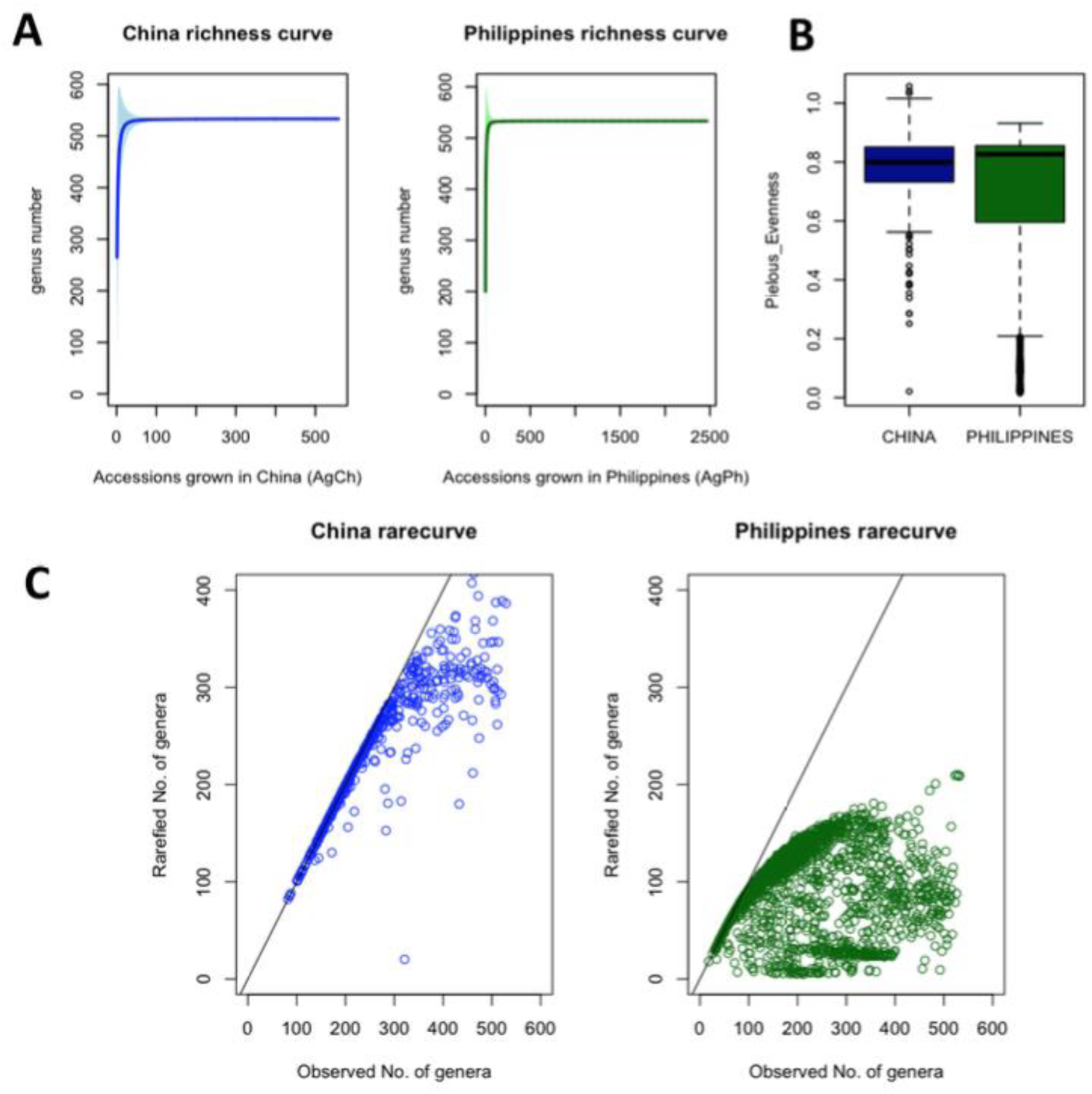
Metagenome sequencing of the 3K-RGP accessions captures leaf microbiome diversity. **A** Richness curves for accessions grown in China and in Philippines. The y-axis represents number of identified genera and x-axis the number of collected samples. The shade on the curves represents the confidence interval of two in the curve points. **B** Evenness bar plots with Pielou’s formula. **C** The rarefaction curves for China and Philippines microbiomes showed the number of expected genera reach a plateau between 100 to 300 observed genera. The line indicates the theoretical linear correlation for rarefaction curves. All 3,024 accessions from the 3K-RGP and the 600 genera found in the analysis were used for the curves. Full data is in Supplementary Table S2.

**FigS3.**
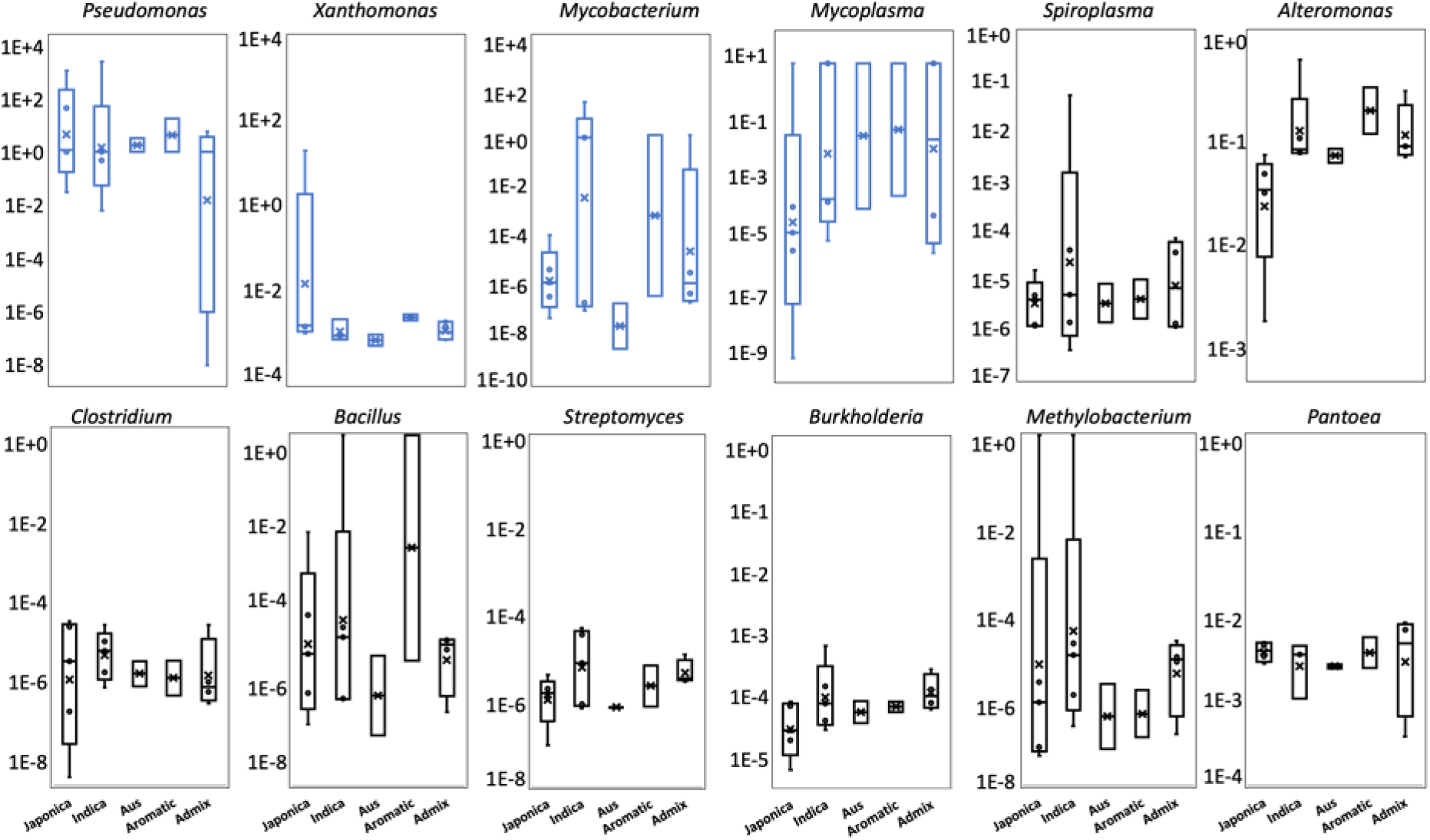
Member of the rice microbial community are present in accessions grown in Philippines. Logarithmic relative abundance of some bacterial groups found in the rice microbiome using specific 16S genus primers. 18 accessions from the 3K-RGP were validated for 12 groups of bacteria present in the rice microbiome. The most abundant bacteria are indicated in blue.

**Fig.S4.**
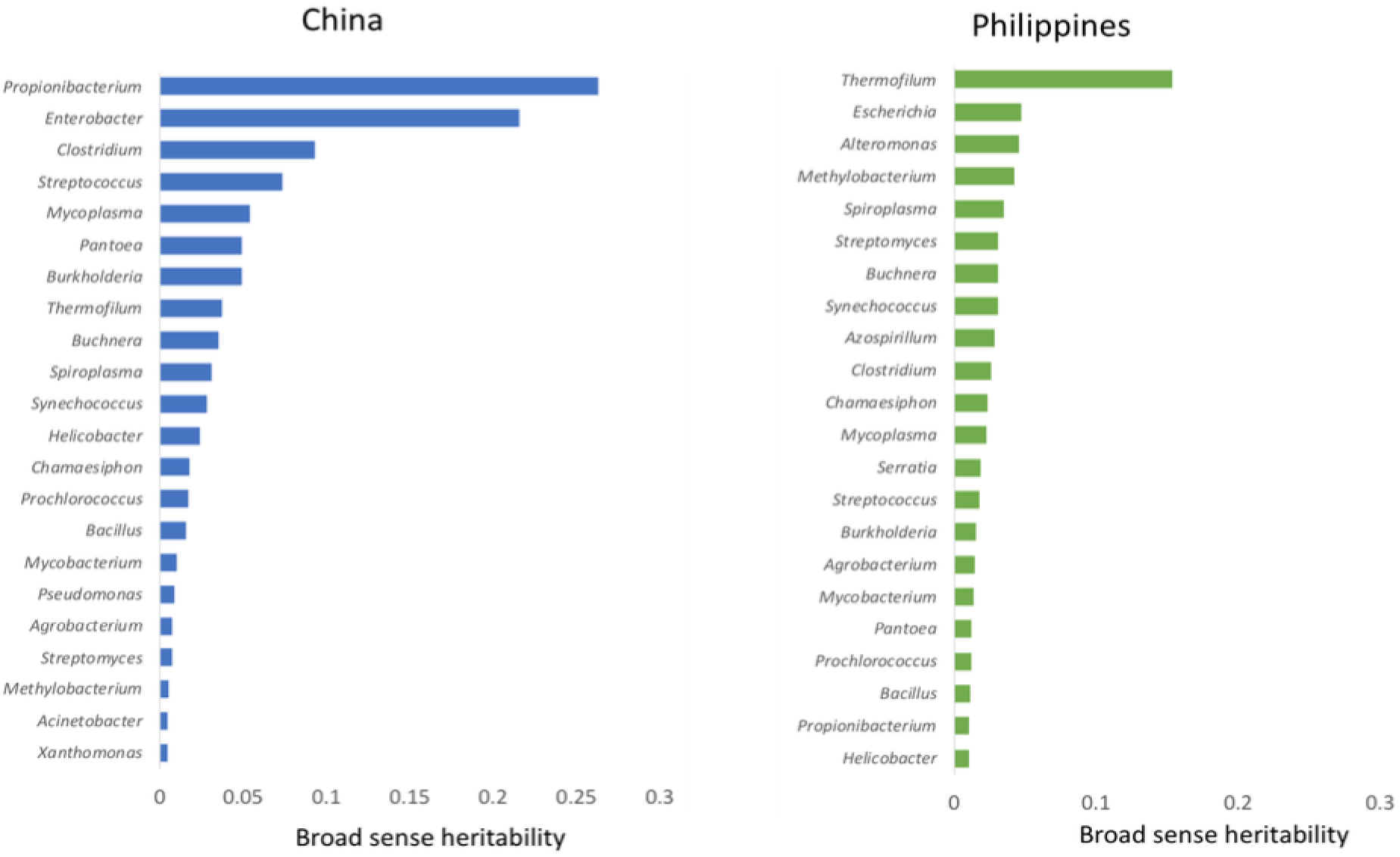
Rice varietal groups associates with variation of few genera. Broad-sense heritability estimates for China and Philippines genera. The heritability was calculated for the most abundant genera in each environment with a random linear model.

**Fig.S5.**
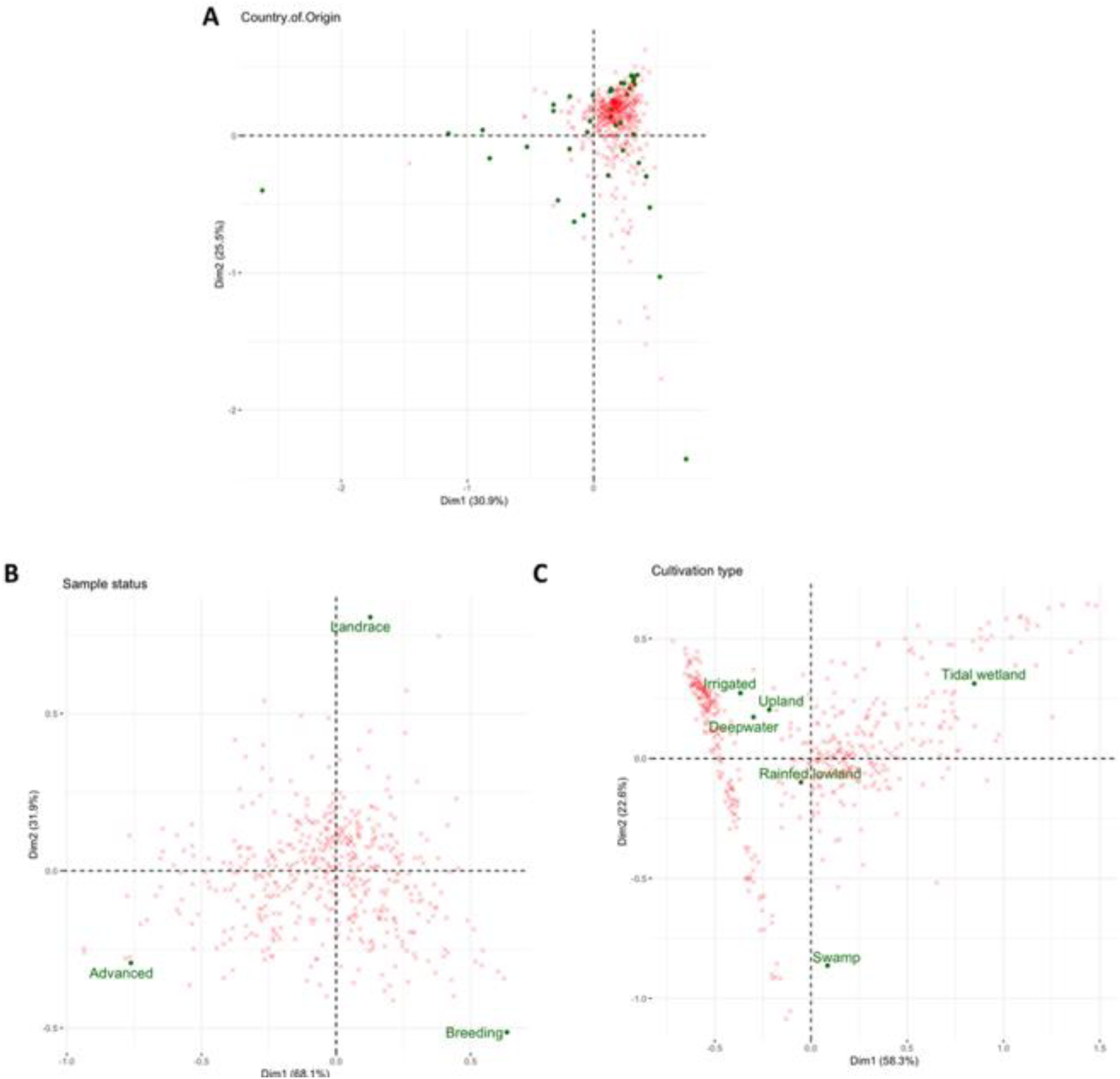
The distribution of the rice leaf microbiome affected by additional factors. Canonical correspondence analysis showing the distribution of microbial genera and rice accessions classified as **A** country of origin, **B** breeding classification, and **C** ecosystem. Red crosses represent the 533 genera found in the leaf microbiome of 467 accessions grown in Philippines. The chi-square values for each plot was 41.7436, P-value < 0.05.

**Fig.S6.**
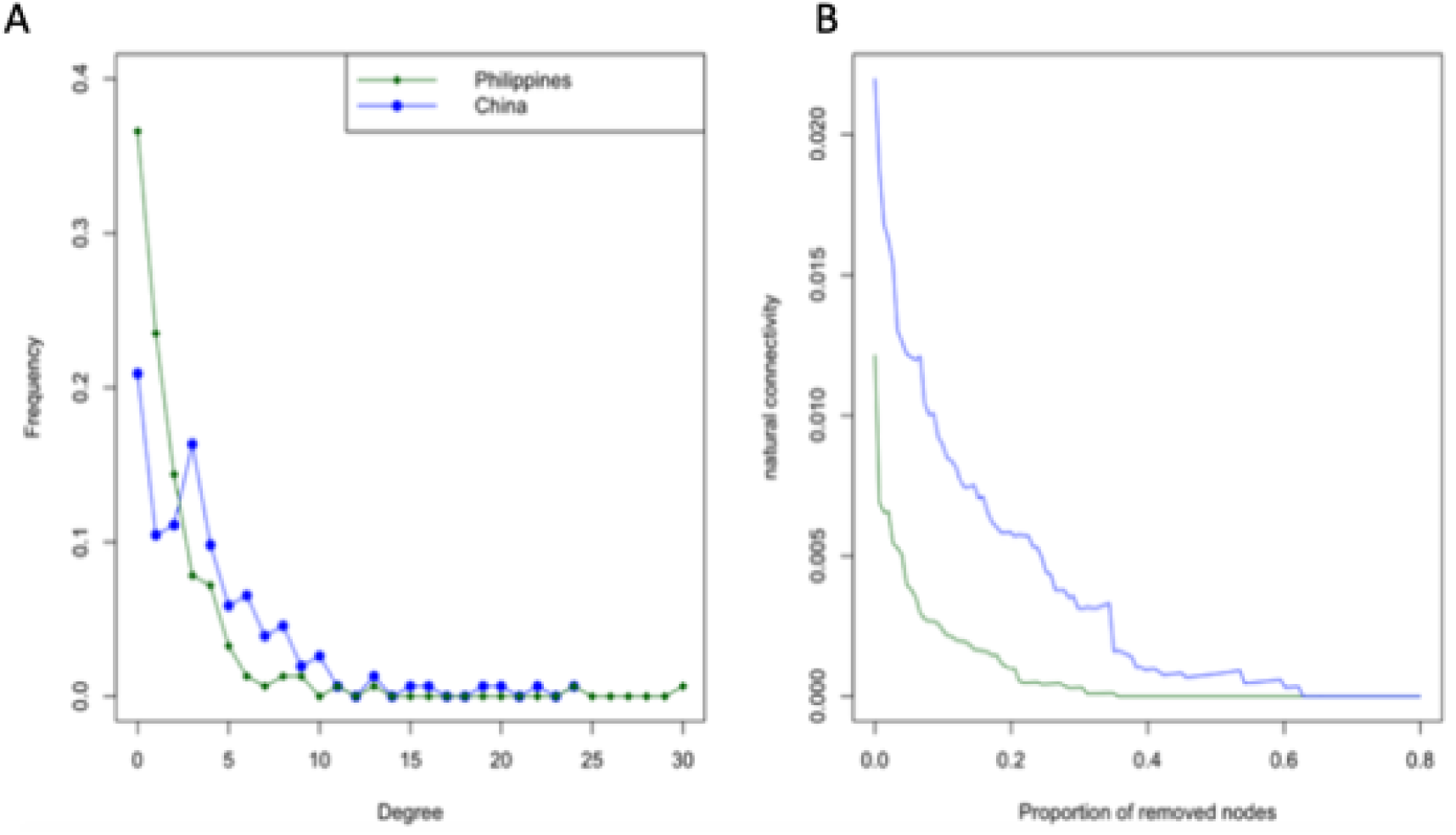
The microbial ecological networks from China and Philippines display similar connectivity and stability features. **A** Frequency of connections (degree) across the network**. B** Network stability plot based on the effect of removing nodes in the network (betweenness centrality) for China (blue) and the Philippines (green).

**Fig.S7.**
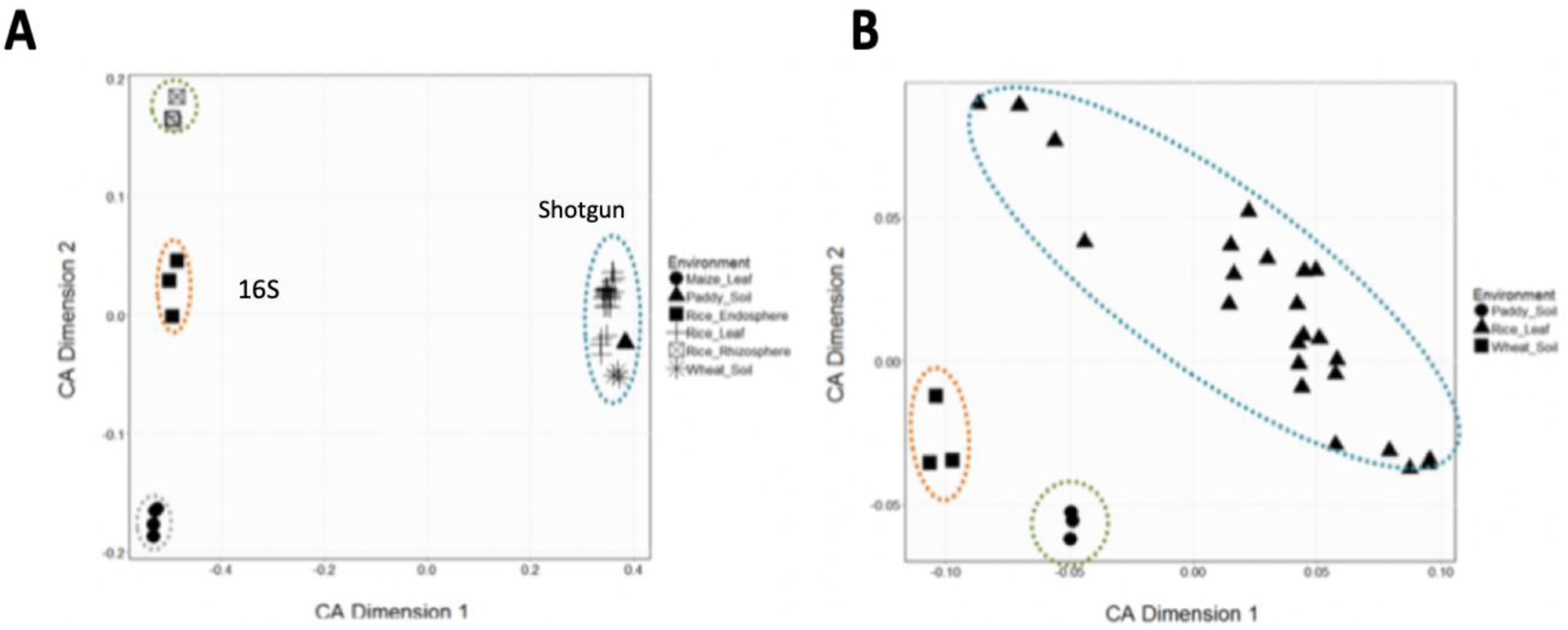
The leaf, roots and soil microbiomes have different functional profiles. Correspondence analysis of functional profiles from different microbiome datasets. **A** Correspondence analysis from databases with 16S amplicon and shotgun reads sequencing approaches. CA Dimension 1 and CA dimension 2 explains 80% and 10% of the differences. **B** Correspondence analysis of databases with shotgun sequences. Dimension 1 and dimension 2 explains 65% and 30% of the differences. We used 19 shotgun databases obtained from NCBI and our dataset. Due to the number of samples for our data compared with the NCBI data, we used median relative abundance of pathways per variety (N=24).

